# FAP-intrinsic Hedgehog signaling controls intramuscular adipogenesis, fibrosis, and myofiber regeneration

**DOI:** 10.64898/2026.05.29.727896

**Authors:** Xinyue Liu, Vincent He, Benedict Deepesh, Alessandra Norris, Daniel Kopinke

## Abstract

Fibro/adipogenic progenitors (FAPs) are multipotent stromal cells that support myofiber regeneration, but can also give rise to intramuscular adipose tissue (IMAT) and fibrotic scar tissue. While the Hedgehog pathway suppresses FAP adipogenesis and promotes myofiber repair through ligand Desert Hedgehog, the key cell type that senses this signal has remained unclear. Here, we demonstrate through FAP-specific deletion of the Hedgehog signal transducer Smoothened that FAPs are the primary Hedgehog-responding cells during muscle regeneration. Loss of Smoothened in FAPs increases IMAT, causes persistent fibrosis, reduces the Hedgehog-dependent effectors TIMP3 and GDF10, and impairs myofiber regeneration. FAPs lacking Smoothened also fail to support *in vitro* myoblast differentiation and fusion as efficiently as control FAPs, showing that Hedgehog signaling helps establish a pro-myogenic FAP state early after injury. Pharmacological Hedgehog activation via the Smoothened agonist SAG fails to rescue adipocyte accumulation or myofiber regeneration when FAPs lack Smoothened. Together, these findings provide direct genetic evidence that FAPs are the primary cellular mediators of Hedgehog signaling in muscle and establish FAP Hedgehog signaling competence as a key determinant of regenerative outcome and a target for restoring muscle repair in disease.

## Introduction

Successful skeletal muscle regeneration depends on coordinated interactions between muscle stem cells (MuSCs) and the surrounding stromal niche. Central to this niche are fibro/adipogenic progenitors (FAPs), a multipotent stromal population that supports MuSC-driven repair through the secretion of pro-myogenic factors (Joe et al. 2010, Lukjanenko et al. 2019, Wosczyna et al. 2019, Uezumi et al. 2021). However, with age and disease, FAPs differentiate into adipocytes and myofibroblasts, giving rise to intramuscular adipose tissue (IMAT) and fibrosis that impair muscle regeneration (Joe et al. 2010, Uezumi et al. 2010, Uezumi et al. 2011, Lemos et al. 2015, Kopinke et al. 2017, Hogarth et al. 2019, Santini et al. 2020). Importantly, IMAT itself directly restricts functional muscle recovery (Biltz et al. 2020, Norris et al. 2025, Palzkill et al. 2026), underscoring that the signals governing FAP fate are directly relevant to regenerative outcomes.

Both FAPs and MuSCs can possess primary cilia, small antenna-like structures located on their cell surface that receive and transmit signals from the environment (Fu et al. 2014, Jaafar Marican et al. 2016, Kopinke et al. 2017, Palla et al. 2022). Among the many signals cilia detect, Hedgehog (Hh) signaling is a key ciliary pathway involved in stem cell fate decisions, tissue maintenance, and regeneration (Anvarian et al. 2019, Kopinke et al. 2021, Ingham 2022). In muscle, our previous work established Desert Hedgehog (DHH) as the key Hh ligand (Kopinke et al. 2017, Norris et al. 2023), which binds the receptor Patched1 (PTCH1) on the cilium, relieving PTCH1-mediated inhibition of the signal transducer Smoothened (SMO) to activate the pathway (Goetz and Anderson 2010, Kopinke et al. 2021). Loss of *Dhh* results in excess IMAT formation and impaired muscle regeneration. Conversely, we and others have shown that global Hh activation via the SMO agonist SAG blocks intramuscular adipocyte accumulation (Norris et al. 2023) and improves muscle regeneration (Palla et al. 2022, Norris et al. 2023). However, because both FAPs and MuSCs possess primary cilia and are, therefore, competent to sense Hh signals, it remains unclear whether this muscle phenotype reflects a direct effect of Hh on MuSCs themselves, or an indirect effect mediated through FAP-derived paracrine signals controlled by Hh.

Here, using FAP-specific genetic deletion of *Smo*, we demonstrate that FAP-intrinsic Hh signaling is the key for suppressing pathological adipose and fibrosis, and for sustaining the pro-myogenic paracrine FAP secretome that supports myofiber regeneration. Specifically, we developed a novel mouse model in which we deleted the Hh effector SMO specifically and only in FAPs (FAP^no SMO^), thereby preventing FAPs from activating Hh signaling even in the presence of the ligand DHH. As a result, FAP^no SMO^ mice showed more IMAT and persistent fibrosis after injury, demonstrating that FAP-intrinsic SMO signaling is required to suppress these pathological fate conversions. Notably, FAP^no SMO^ mice also exhibited smaller, poorly regenerated myofibers, pointing to an indirect requirement for Hh in muscle repair, one mediated through the FAP paracrine secretome rather than through direct Hh action on MuSCs. Importantly, pharmacological activation of Hh via the Smoothened agonist, SAG, failed to rescue the FAP^no SMO^ phenotype, confirming that FAPs are the direct responders of Hh during muscle regeneration, and that Hh promotes muscle repair indirectly, by acting on FAPs to sustain a pro-myogenic secretome that supports muscle stem cell-driven regeneration. Together, our results establish FAPs as the core cellular mediators through which Hh limits IMAT and fibrosis, and point to FAP-specific Hh signaling as a target for restoring efficient muscle regeneration in disease.

## Results

### FAP-specific *Smo* deletion abolishes ciliary SMO accumulation and Hh target gene expression

To directly manipulate Hh pathway specifically in FAPs, we generated mice carrying a FAP-specific conditional deletion of *Smo* by crossing *Pdgfrα*^*CreERT*^ mice with *Smo*^*fl/fl*^ mice (Fig. 1A). PDGFRα is the gold standard marker to identify FAPs (Joe et al. 2010, Uezumi et al. 2010, Uezumi et al. 2011, Uezumi et al. 2014, Kopinke et al. 2017, Hogarth et al. 2019, Lukjanenko et al. 2019, Wosczyna et al. 2019, Santini et al. 2020). We then injured the tibialis anterior (TA) muscle with cardiotoxin (CTX), a snake venom-derived myotoxin that causes synchronous myofiber necrosis and robustly induces Hh signaling via upregulation of its ligand DHH (Norris et al. 2023), and collected tissue at 3, 7, or 21 days post-injection (dpi) (Fig. 1B).

**Figure 1.**
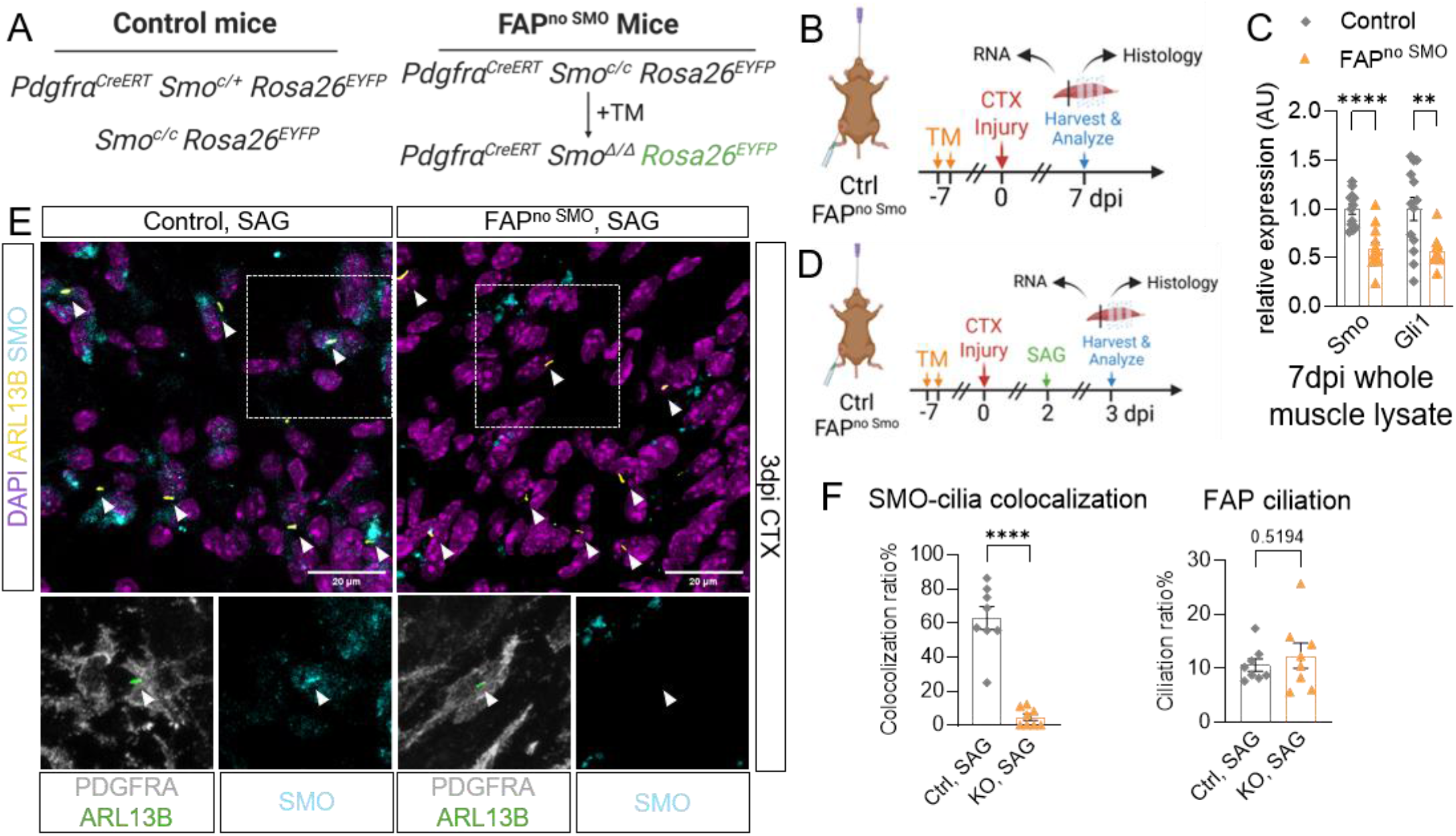
FAP-specific *Smo* deletion blocks Hh signaling and eliminates ciliary SMO accumulation. **(A)** Genetic scheme illustrating the *Pdgfrα*^*CreERT*^; *Smo*^fl/fl^ conditional knockout strategy. **(B)** Experimental timeline for cardiotoxin (CTX)-induced tibialis anterior (TA) muscle injury. **(C)** RT-qPCR analysis of *Smo* and *Gli1* transcript levels in TA whole-muscle lysates (n = 9–10 mice per group) from control and FAP^no SMO^ mice at 7 days post-CTX injection (dpi). Data are presented as mean ± SEM. **(D)** Experimental outline. SAG was administered at 2 dpi, and TA cross sections were collected at 3 dpi. **(E)** Representative immunofluorescence images, and **(F)** quantification of FAP ciliation and SMO–cilia colocalization (normalized to FAP number; n = 5–6 mice per group) in SAG-treated control and FAP^no SMO^ TA cross sections at 3 dpi. Top row: DAPI (purple), nuclei; ARL13B (yellow), cilia; SMO (cyan). Scale bar, 20 µm. Bottom row (zoom-in): PDGFRα (gray); ARL13B (green), cilia; SMO (cyan). ****p < 0.0001; **p < 0.01. Statistical comparisons were made by unpaired two-tailed Student’s t-test (C, F).

To confirm conditional deletion, we performed RT-qPCR on the whole TA muscle at 7 dpi. FAP^no SMO^ mice had significantly lower *Smo* levels and a large drop in *Gli1* (GLI Family Zinc Finger 1) expression, a direct readout of active Hh signaling (Kopinke et al. 2017, Norris et al. 2023), confirming that loss of SMO in FAPs effectively turns off Hh signaling (Fig. 1C). Because binding of DHH to PTCH1 relieves PTCH1-mediated inhibition of SMO, allowing SMO to enter and accumulate within the primary cilium (Goetz and Anderson 2010, Kopinke et al. 2021), we next asked whether SMO protein was correspondingly absent from FAP cilia in FAP^no SMO^ mice. To maximally activate the pathway, we administered SAG on day 2 post-CTX injury and examined TA cross sections at 3 dpi by immunofluorescence (Fig. 1D). In control mice, SAG strongly promoted SMO accumulation in cilia (ARL13B^+^) on FAPs (PDGFRα^+^) (Fig. 1E). In FAP^no SMO^ mice, SMO was almost completely absent from cilia, despite unchanged FAP ciliation ratios (Fig. 1F). These results validate the FAP^no SMO^ model and confirm that SMO is needed for Hh signal transduction within FAP cilia after injury.

### FAP-specific SMO deletion promotes intramuscular adipogenesis

In normal muscle repair, FAPs expand transiently after injury and are subsequently cleared (Lemos et al. 2015), with only approximately 20% differentiating into adipocytes (Norris et al. 2025). We previously showed that loss of DHH results in increased IMAT formation after acute injury (Norris et al. 2023). Therefore, we asked whether FAP-specific loss of SMO phenocopies this global Hh loss-of-function state, driving excess IMAT accumulation *in vivo*. For this, we injured the TA muscle of FAP^no SMO^ and control mice with CTX, and collected TA muscle at 21 dpi (Fig. 2A). To quantify adipocyte accumulation, we stained TA sections against PERILIPIN, a lipid droplet-coating protein and marker of mature adipocytes, at 21 dpi (Fig. 2A). FAP^no SMO^ mice displayed a robust and significant increase in PERILIPIN^+^ adipocytes compared to controls, consistent across both sexes (Fig. 2A), demonstrating that DHH is sensed by FAPs to suppress their adipogenic differentiation, and phenocopies the IMAT excess observed in *Dhh*^−*/*−^ mice (Norris et al. 2023).

**Figure 2.**
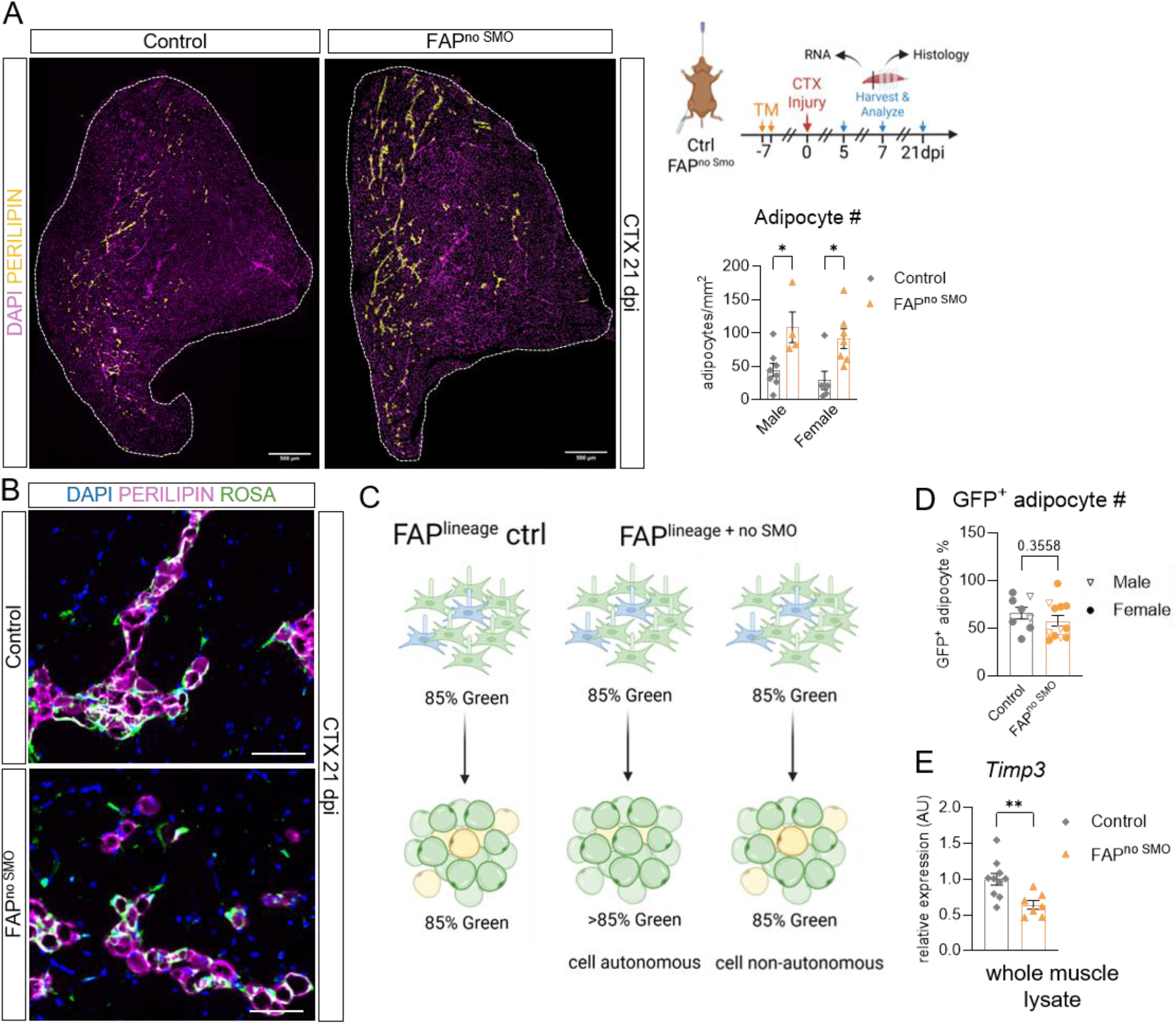
FAP-specific SMO deletion promotes intramuscular adipogenesis. **(A)** Experimental outline (right top), representative immunofluorescence images and adipocyte quantification of TA cross sections from control and FAP^no SMO^ mice at 21 dpi CTX. Left: tile-scan images stained for DAPI (purple), nuclei, and PERILIPIN (yellow), adipocytes. Scale bar, 500 µm. Symbols indicate sex. Right bottom: quantification of intramuscular adipocyte density from TA cross sections at 21 dpi CTX. Adipocytes per mm^2^ of injured area; n = 4–8 mice per group per sex. **(B)** Representative zoom-in immunofluorescence images of TA cross sections from control and FAP^no SMO^ mice at 21 dpi CTX. DAPI (blue), nuclei; PERILIPIN (purple), adipocytes; Rosa26-GFP (green), FAP-derived cells. Scale bar, 50 µm. **(C)** Model of cellular effect for adipogenesis. The percentage of EYFP^+^ adipocytes (green) would argue for a cell-autonomous effect after loss of FAP SMO, whereas an increase in total number of both EYFP^+^ and EYFP^−^ adipocytes (without affecting the proportion expressing EYFP) would suggest a non-cell-autonomous mechanism. **(D)** Percentage of lineage-traced (GFP^+^) adipocytes among total PERILIPIN^+^ adipocytes (n = 8–9 mice per group). Symbols indicate sex. **(E)** RT-qPCR analysis of *Timp3* in TA whole-muscle lysates (n = 6–10 mice per group) from control and FAP^no SMO^ mice at 7 dpi CTX. Data are presented as mean ± SEM. **p < 0.01; *p < 0.05. Statistical comparisons were made by unpaired two-tailed Student’s t-test (A, D, E).

To determine whether this represented a cell-autonomous or non-cell-autonomous effect, we used a Rosa26-EYFP lineage tracing approach to permanently label the cells that derived from the FAPs with successful removal of SMO (Fig. 1A, 2B) (Srinivas et al. 2001, Kopinke et al. 2017). Since the recombination efficiency of the *Pdgfrα*^*CreERT/+*^ allele achieves ∼80–85% recombination efficiency (Norris et al. 2025), approximately 15–20% of FAPs retain an intact *Smo* allele and will therefore remain GFP^-^ after tamoxifen administration. As endogenous EYFP fluorescence is diminished by tissue fixation, lineage-traced cells were detected using an anti-GFP antibody to amplify the signal. If SMO loss promotes cell-autonomous adipogenesis of FAPs, we would expect a larger fraction of adipocytes to be GFP^+^ in FAP^no SMO^ mice (Fig. 2C). Instead, the proportion of PERILIPIN^+^ adipocytes that were also GFP^+^ was similar between FAP^no SMO^ and control mice (Fig. 2D), despite more total adipocytes overall. The comparable labeling proportions indicate that both GFP^+^ and GFP^−^ FAPs contribute equally to the expanded adipocyte pool, suggesting that FAP SMO loss promotes adipogenesis through a non-cell-autonomous mechanism rather than through cell-autonomous differentiation (Fig. 2C). This is consistent with our previous finding, where we found that Hh-responding FAPs suppress the adipogenic differentiation of neighboring FAPs through a paracrine mechanism (Kopinke et al. 2017).

We previously identified TIMP3 (Tissue Inhibitor of Metalloproteinase 3), a secreted metalloproteinase inhibitor and direct Hh target gene in FAPs that suppresses adipogenesis by blocking MMP14 (Matrix Metalloproteinase 14), a membrane-anchored protease that promotes adipogenic extracellular matrix remodeling (Kopinke et al. 2017, Norris et al. 2023). To determine if *Timp3* was impacted and involved in this adipogenic suppression, we performed RT-qPCR on whole TA lysates at 7 dpi and revealed significantly reduced *Timp3* expression in FAP^no SMO^ mice compared to controls (Fig. 2E), confirming that FAP-intrinsic Hh signaling is required to maintain *Timp3*-mediated suppression of adipogenesis *in vivo*. Together, these data establish that Hh pathway activity within FAPs is required to maintain *Timp3*-dependent suppression of adipogenesis, and that its loss causes excess adipogenesis through a non-cell-autonomous mechanism.

### Absence of Hh activity in FAPs drives persistent intramuscular fibrosis

Besides adipocytes, FAPs can also differentiate into myofibroblasts, which contribute to fibrotic scar tissue formation (Uezumi et al. 2011, Lemos et al. 2015). This pathological dual fate conversion is a hallmark of conditions including muscular dystrophy, aging, and chronic limb-threatening ischemia (CLTI), where IMAT and fibrosis accumulate concurrently in muscle (Hogarth et al. 2019, Lukjanenko et al. 2019, Uezumi et al. 2021, Palzkill et al. 2026). To determine whether removing the ability of FAPs to respond to DHH also affects myofibroblast differentiation, we investigated whether FAP-specific SMO loss also promoted a fibrogenic fate shift (Fig. 3A). Sirius red staining of TA cross sections revealed a significant increase in fibrotic area in FAP^no SMO^ mice compared to controls at both 7 and 21 dpi, consistent across sexes (Fig. 3A and Supplementary Fig. 1). These findings demonstrate that FAP-specific loss of Hh signaling drives persistent muscle fibrosis in both sexes.

**Figure 3.**
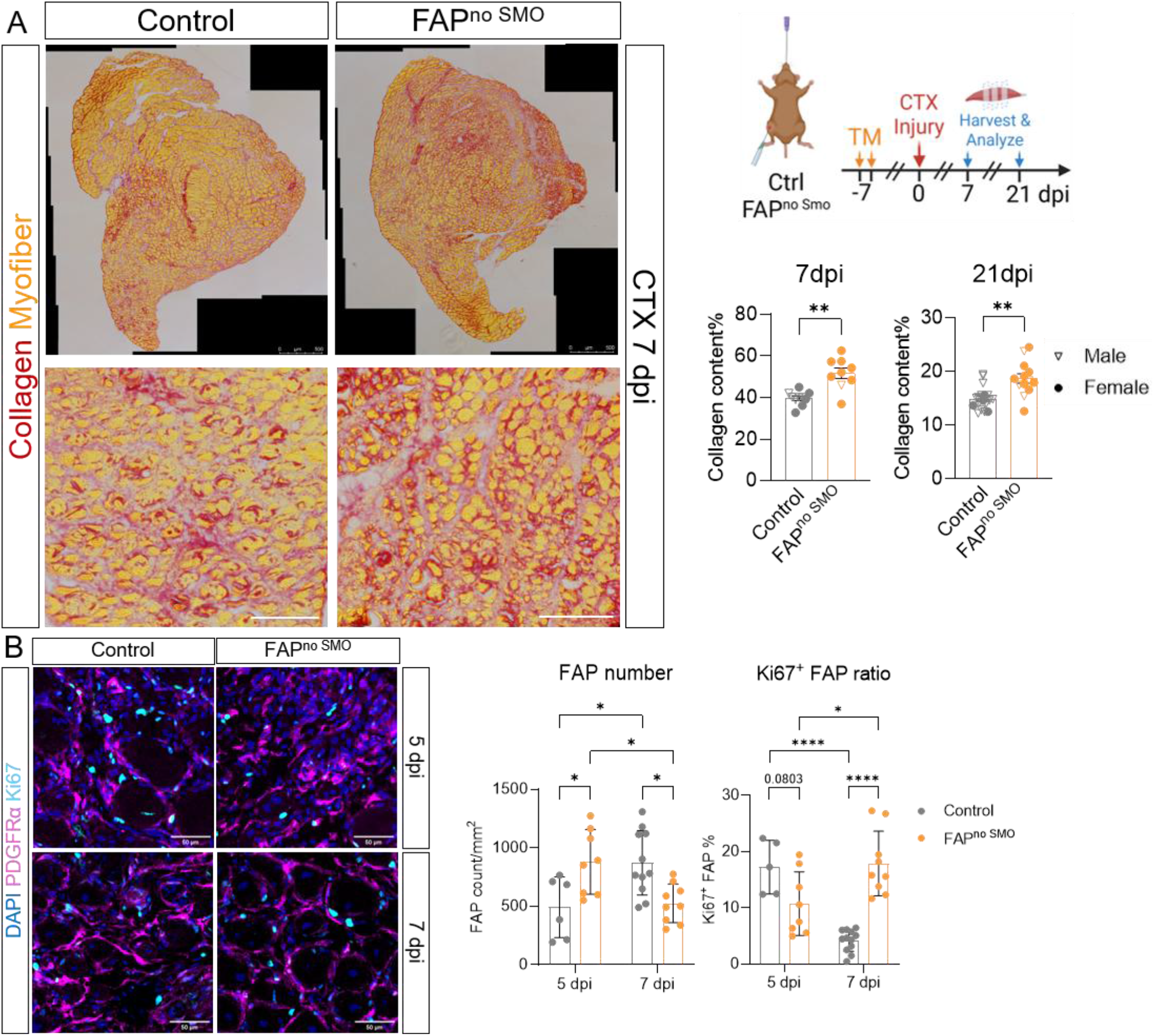
Hedgehog signaling deficiency in FAPs alters FAP proliferation dynamics and drives persistent muscle fibrosis. **(A)** Experimental outline (right top), representative Sirius red-stained images of TA cross sections from control and FAP^no SMO^ mice at 7 dpi CTX showing whole-section (top; scale bar, 500 µm) and zoom-in (bottom; scale bar, 300 µm) views; collagen, red; myofibers, yellow. Right bottom: quantification of fibrotic area (percentage of injured area) at 7 dpi (n = 9–10 mice per group) and 21 dpi CTX (n = 9 mice per group); symbols indicate sex. **(B)** Left: representative immunofluorescence of TA cross sections stained for Ki67 (cyan), PDGFRα (purple), and DAPI (blue) at 5 and 7 dpi CTX. Scale bar, 50 µm. Center: quantification of total PDGFRα^+^ FAP number per whole TA cross section. Right: percentage of proliferating FAPs (Ki67^+^PDGFRα^+^ cells as a fraction of all FAPs) at 5 and 7 dpi in control and FAP^no SMO^ mice (n = 7–9 mice per group). Data are presented as mean ± SEM. ****p < 0.0001; **p < 0.01; *p < 0.05. Statistical comparisons were made by unpaired two-tailed Student’s t-test (A) or two-way ANOVA with Tukey’s post-hoc test (B).

Since FAPs are the cellular origin of both adipose and fibrotic scar tissue (Joe et al. 2010, Uezumi et al. 2010, Uezumi et al. 2011), we asked whether the concurrent increase in adipocytes and/or collagen deposition in FAP^no SMO^ mice reflects an expanded FAP progenitor pool driven by excess proliferation. To address this, we immunostained TA cross sections for Ki67 (a nuclear marker of actively cycling cells) and PDGFRα to quantify proliferating FAPs at 5 and 7 dpi (Fig. 3B). In control mice, FAP numbers increased between 5 and 7 dpi, accompanied by a progressive decline in the proportion of Ki67^+^ FAPs, consistent with the normal pattern of transient expansion followed by cell-cycle exit and clearance (Joe et al. 2010, Uezumi et al. 2010, Lemos et al. 2015, Theret et al. 2021). In FAP^no SMO^ mice, this dynamic was disrupted that at 7 dpi, when wild-type FAPs are exiting the cell cycle, FAP^no SMO^ mice retained a markedly higher fraction of Ki67^+^ FAPs alongside a greater total number of PDGFRα^+^ cells, indicating that FAPs fail to exit the cell cycle in a timely manner upon loss of SMO. Since intramuscular adipogenesis is initiated as early as day 3 post-injury (Norris et al. 2025), before the FAP proliferative window we observed at days 5 and 7, FAPs that are still cycling at 7 dpi are unlikely to be entering the adipogenic lineage. Instead, this late proliferative phase, beyond the normal adipogenic commitment window, coincides with the accumulation of fibrotic collagen deposition (Lemos et al. 2015). We therefore propose that this expanded pool of late-cycling FAPs preferentially feeds the fibrogenic rather than the adipogenic lineage, providing a cellular basis for the excess fibrosis observed in FAP^no SMO^ mice.

### Loss of FAP Hh signaling impairs myofiber regeneration

FAPs support MuSC-driven muscle repair through secreting pro-myogenic factors (Lukjanenko et al. 2019, Wosczyna et al. 2019). To test whether the myofiber regeneration defects observed in *Dhh* loss-of-function mice (Norris et al. 2023) reflect a disruption of Hh-controlled FAP-specific myogenic factors, we measured myofiber cross-sectional area (CSA) in TA sections stained with fluorophore-conjugated phalloidin, which labels filamentous actin and delineates individual myofiber boundaries, at both 7 and 21 dpi (Fig. 4A).

**Figure 4.**
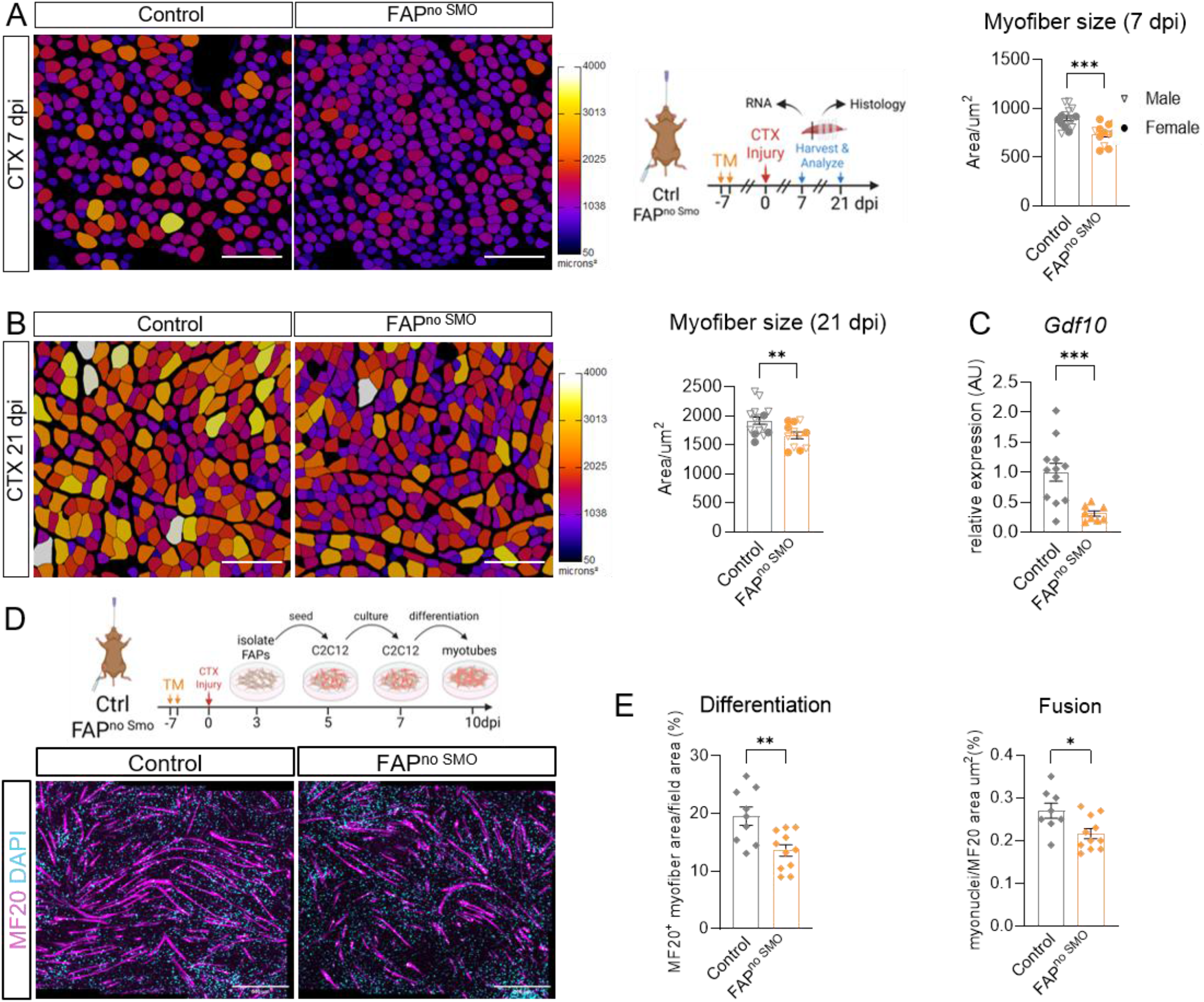
Loss of Hedgehog signaling in FAPs impairs myofiber regeneration both *in vivo* and *in vitro*. **(A)** Experimental outline (middle), representative images and quantifications of average myofiber cross-sectional area (CSA) at 7 dpi CTX in control and FAP^no SMO^ mice (n = 10–15 mice per group). Left: representative images of TA cross sections from control and FAP^no SMO^ mice at 7 dpi. Individual myofibers were immunolabeled with phalloidin and pseudo-color-coded by CSA. Scale bar, 200 µm. Right: average CSA per animal; symbols indicate sex. **(B)** Representative images and quantifications of average myofiber CSA at 21 dpi CTX in control and FAP^no SMO^ mice (n = 5–8 mice per sex per group). Left: representative images of TA cross sections from control and FAP^no SMO^ mice at 21 dpi. Individual myofibers were immunolabeled with phalloidin and pseudo-color-coded by CSA. Scale bar, 200 µm. Right: average CSA per animal; symbols indicate sex. **(C)** RT-qPCR analysis of *Gdf10* in TA whole-muscle lysates (n = 7–10 mice per group) from control and FAP^no SMO^ mice at 7 dpi CTX. Data are presented as mean ± SEM. **(D)** Schematic of the coculture experimental design (top). FAPs isolated from control and FAP^no SMO^ mice were cocultured with C2C12 myoblasts in C2C12 differentiation media. Representative immunofluorescence images of C2C12-derived myotubes at 3 days post-myogenic induction (bottom). MF20 (myosin heavy chain; purple), mature myofibers; DAPI (cyan), nuclei. Scale bars: 500 µm. **(E)** Quantification of the differentiation index (MF20^+^ area as percentage of total field area) and fusion index (nuclei per MF20^+^ area) of C2C12-derived myotubes (n = 9–12 replicates per group). Data are represented as mean ± SEM. ***p < 0.001; **p < 0.01; *p < 0.05. Statistical comparisons were made by unpaired two-tailed Student’s t-test (B–F).

Using our automated segmentation pipeline to quantify myofiber size from phalloidin-stained sections pseudo-color-coded by CSA (Waisman et al. 2021), we found that at 7 dpi, FAP^no SMO^ muscles had significantly smaller myofibers than controls, indicating impaired early myofiber regeneration (Fig. 4A and Supplementary Fig. 2). At 21 dpi, FAP^no SMO^ muscles continued to show lower average fiber CSA and this deficit was observed consistently in both male and female mice (Fig. 4B and Supplementary Fig. 2). We have previously shown that Hh signaling in FAPs induces GDF10 (Growth Differentiation Factor 10), a secreted pro-myogenic factor that promotes myofiber growth (Uezumi et al. 2021, Norris et al. 2023). To test whether the myofiber regeneration defects in FAP^no SMO^ mice reflect a failure to induce GDF10 due the inability of FAPs to activate Hh signaling, we performed RT-qPCR on whole TA lysates at 7 dpi and found that *Gdf10* was indeed significantly reduced in FAP^no SMO^ mice (Fig. 4C). This data demonstrates that FAP-intrinsic Hh activity is required not only to restrain adipogenesis, but also to support myofiber regeneration through paracrine mechanisms.

To directly test whether this FAP paracrine support requires FAP-intrinsic Hh signaling, we employed a coculture assay *in vitro* (Fig. 4D). FAPs isolated from FAP^no SMO^ or control mice were cocultured with C2C12 myoblasts in differentiation media. After 3 days of myogenic induction, cultures were stained for MF20 (myosin heavy chain), which marks terminally differentiated myotubes, and DAPI to label all nuclei (Fig. 4D). Compared to control FAPs, FAPs without SMO significantly reduced both the differentiation index (MF20^+^ myofiber area as a percentage of total field area) and the fusion index (myonuclei per MF20^+^ area) of C2C12-derived myotubes (Fig. 4D, 4E). Critically, this coculture system contains no exogenous Hh ligand. FAP^no SMO^ FAPs therefore cannot transduce a Hh signal *in vitro* regardless of treatment. The impaired myogenic support thus reflects an intrinsically altered secretory state that FAP^no SMO^ cells carry over from *ex vivo* into the co-culture. This indicates a cell-autonomous property of SMO deficiency, instead of a failure to respond to ligands, which are absent from the dish.

Together, these findings demonstrate that Hh signaling in FAPs shapes the FAP secretome in a cell-autonomous manner, and that this paracrine program that supports muscle formation, complementing the fiber regeneration defects seen in FAP^no SMO^ mice, and is indispensable for productive myoblast differentiation and fusion (Joe et al. 2010, Lukjanenko et al. 2019, Norris et al. 2023).

### FAP-specific *Smo* deletion blocks responsiveness to pharmacological Hh activation

To determine whether FAPs require SMO to respond to pharmacological Hh activation *in vivo*, we treated control and FAP^no SMO^ mice with SMO agonist SAG on days 0, 2, and 4 post-CTX injury and harvested TA muscles at 7 dpi (Fig. 5A). In control mice, SAG robustly increased expression of the canonical Hh target genes *Gli1* and *Ptch1*, as well as the Hh-dependent FAP effector genes *Gdf10* (pro-myogenic) and *Timp3* (anti-adipogenic) (Fig. 5B). Importantly, this response was significantly blunted or absent in FAP^no SMO^ mice. Thus, loss of SMO renders FAPs unable to respond to exogenous Hh pathway activation, even when SAG is delivered systemically.

**Figure 5.**
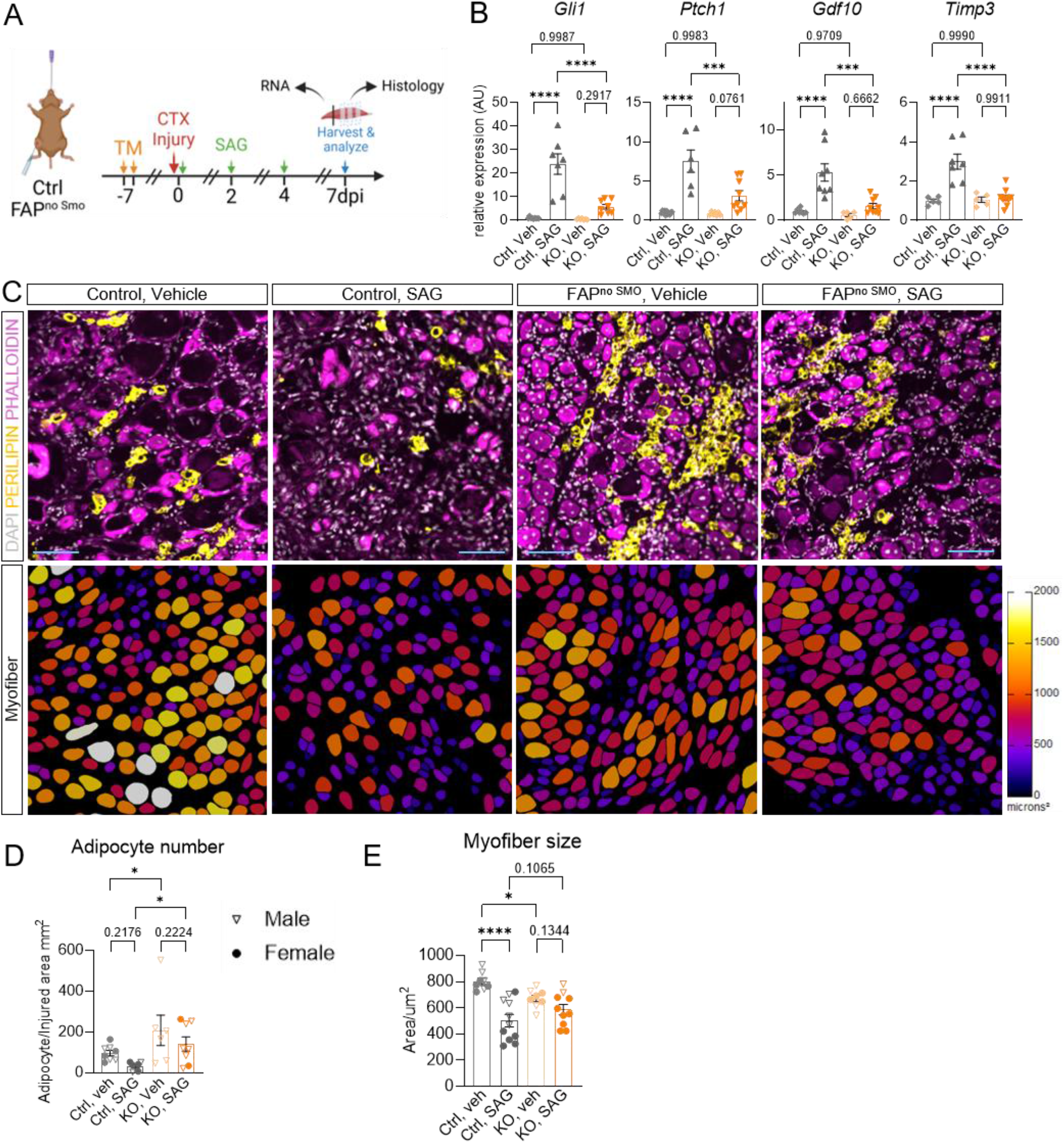
FAP-specific *Smo* deletion blocks transcriptional and tissue responses to SAG. (**A)** Experimental scheme for SAG treatment (intraperitoneal injections on days 0, 2, and 4 post-CTX injury); tissue harvested at 7 dpi. **(B)** RT-qPCR analysis of *Gli1, Ptch1, Gdf10*, and *Timp3* in TA whole-muscle lysates (n = 6–8 mice per group) from control vehicle, control SAG, FAP^no SMO^ vehicle, and FAP^no SMO^ SAG mice at 7 dpi CTX. **(C)** Representative immunofluorescence (top) and pseudo-color-coded (bottom) images of TA cross sections across treatment groups at 7 dpi CTX. Top: DAPI (gray), nuclei; PERILIPIN (yellow), adipocytes; phalloidin (purple), myofibers. Scale bar, 60 µm. Bottom: myofibers pseudo-color-coded by CSA. **(D)** Quantification of adipocyte density (adipocytes per mm^2^ of injured area) from TA cross sections at 7 dpi CTX (n = 8–9 mice per group). **(E)** Average myofiber CSA from TA cross sections at 7 dpi CTX (n = 6–8 mice per group). Symbols indicate sex. Data are presented as mean ± SEM. ****p < 0.0001; ***p < 0.001; *p < 0.05. Statistical comparisons were made by one-way ANOVA with Tukey’s post-hoc test (B, D, E).

We next asked whether this loss of transcriptional responsiveness was reflected at the tissue level. Consistent with its known anti-adipogenic effect (Kopinke et al. 2017, Norris et al. 2023), SAG reduced PERLIPIN^+^ adipocyte accumulation in control muscles (Fig. 5C, 5D). In contrast, SAG failed to reduce adipocyte accumulation in FAP^no SMO^ muscles, demonstrating that the anti-adipogenic effect of SAG requires FAP-intrinsic SMO-dependent Hh signaling. We then examined myofiber regeneration by quantifying myofiber CSA. Under a repeated dosing regimen, SAG reduced average myofiber size in control muscles, consistent with our previous observation that sustained Hh activation can impair myofiber regeneration (Norris et al. 2023). Importantly, SAG did not improve myofiber size in FAP^no SMO^ mice and did not shift the phenotype toward that of SAG-treated controls (Fig. 5C, 5E). Together, these data show that systemic SAG acts through SMO-dependent Hh signaling in FAPs. When FAPs lack SMO, pharmacological Hh activation no longer induces the Hh-dependent FAP transcriptional program and fails to modify either adipocyte accumulation or myofiber regeneration.

## Discussion

Hedgehog signaling has long been implicated in skeletal muscle regeneration, but the identity of the key cell type sensing and transducing this signal has remained unresolved. Our prior work provided compelling indirect support for FAPs as the primary Hh-responding population by showing that: (a) ablation of FAP cilia suppressed IMAT formation via Hh de-repression and low-level FAP-intrinsic Hh activation (Kopinke et al. 2017), (b) FAPs, not MuSCs, mount a robust transcriptional response to DHH after injury (Norris et al. 2023), and (c) ectopic Hh activation, via FAP-specific removal of *Ptch1*, specifically in FAPs drives fibrosis (Norris et al. 2023). Here, using FAP-specific deletion of *Smo*, we provide direct genetic evidence that FAPs are indeed the key cellular transducers of Hh signaling in skeletal muscle. Loss of SMO in FAPs blocks canonical Hh activation, increases IMAT, promotes persistent fibrosis, and impairs myofiber regeneration post injury. Importantly, systemic pharmacological Hh activation fails to rescue these phenotypes when FAPs lack *Smo*, demonstrating that the relevant regenerative and anti-adipogenic effects of Hh require FAP-intrinsic SMO-dependent Hh signaling.

### FAPs, not MuSCs, are the primary Hh responders during muscle regeneration

Both FAPs and MuSCs possess primary cilia (Fu et al. 2014, Jaafar Marican et al. 2016, Kopinke et al. 2017, Palla et al. 2022) and could, in principle, respond to Hh ligands. However, our data support a model in which FAPs are the main Hh-responding cell type during muscle regeneration. As we show here, FAP-specific loss of *Smo*, which prevents FAPs from responding to Hh ligands, causes the key features seen after loss of the ligand DHH, including increased IMAT and impaired myofiber regeneration (Norris et al. 2023). These findings fit with our previous work showing that FAPs mount a strong transcriptional response to SAG, which is lost upon loss of DHH, whereas MuSCs do not show a robust Hh response when the pathway is activated by SAG or reduced by loss of DHH (Norris et al. 2023). Thus, while MuSCs contain cilia, they may not use them primarily to respond to Hh ligands. Instead, the MuSC loss of cilia phenotype may reflect loss of cilia-dependent GLI3 repression (Palla et al. 2022). For example, loss of MuSC cilia causes defects in quiescence and regeneration (Palla et al. 2022), and *Gli3* deletion causes a related phenotype, with MuSCs leaving quiescence and entering a more activated G_Alert_-like state (Brun et al. 2022). Thus, MuSC cilia may act mainly as guardians of the repressive GLI3 state, whereas FAP cilia both maintain repression and allow ligand-induced SMO activation. Under this interpretation, the failure of SAG to fully rescue force production in mice lacking MuSC cilia (Palla et al. 2022) may reflect a compounded deficit caused by loss of GLI3-repressor-dependent MuSC pool maintenance, rather than a direct requirement for SMO-dependent Hh signaling in MuSCs. We propose that SAG’s pro-regenerative effects in that study are instead mediated primarily through FAPs, which retain intact cilia and SMO and respond robustly to Hh activation by secreting pro-myogenic factors such as GDF10 (Norris et al. 2023). Taken together, the data presented here provide direct genetic proof that FAPs are the primary cellular mediator of Hh signaling in injured muscle.

### The FAP paracrine secretome mediates Hh-dependent myogenic support

The impaired myofiber regeneration in FAP^no SMO^ mice suggests that Hh promotes muscle repair indirectly, by keeping FAPs in a pro-myogenic state. This fits with our previous work showing that Hh induces GDF10 in FAPs and that GDF10 supports myofiber growth (Uezumi et al. 2021, Norris et al. 2023). Here, we found that *Gdf10* expression is reduced in FAP^no SMO^ muscle, suggesting that FAPs need SMO to activate this pro-myogenic program after injury. Our co-culture experiment supports this idea. FAPs isolated from injured FAP^no SMO^ muscle at 3 dpi were less able to support C2C12 differentiation and fusion than control FAPs (Fig. 4D, 4E). Thus, the defect is not only present in the injured muscle, but remains with the FAPs after isolation. This suggests that Hh acts early after injury to help establish a FAP state that supports myogenesis. At the same time, reduced *Timp3* likely explains the increased adipogenesis, since TIMP3 is an Hh-induced anti-adipogenic secreted factor (Kopinke et al. 2017, Norris et al. 2023). Together, our data demonstrate that FAP-intrinsic Hh signaling controls two related repair programs. It limits FAP-to-fat conversion through TIMP3, and it promotes myofiber regeneration by inducing a pro-myogenic FAP program that includes GDF10.

### FAP-intrinsic Hh signaling must be tightly controlled to prevent fatty fibrosis

One unexpected finding is that FAP-specific *Smo* deletion causes persistent fibrosis, a phenotype we did not observe after global loss of DHH. This was surprising because excessive Hh activation, either through FAP-specific Ptch1 deletion or repeated SAG administration, can also drive fibrosis (Norris et al. 2023). Together, these findings suggest that FAP fate depends on a tightly controlled range of Hh activity. Too little or too much pathway activity can disrupt normal repair, but likely through different mechanisms. Most of our data fits a canonical SMO-GLI mechanism, since loss of FAP SMO reduces *Gli1* together with the Hh-dependent FAP effector genes *Timp3* and *Gdf10* (Fig. 2E, 4C). This provides a clear explanation for the increased adipogenesis and impaired myofiber regeneration. However, the fibrosis and FAP proliferation phenotypes (Fig. 3 and Supplementary Fig. 1) distinguish FAP^no SMO^ mice from *Dhh*^*-/-*^ mice, suggesting that additional mechanisms may contribute. One possibility is that SMO deletion blocks signaling downstream of all Hh ligands, not only DHH. However, we did not find any evidence of compensatory upregulation of either SHH or IHH (Kopinke et al. 2017, Norris et al. 2023). Another possibility is that SMO controls aspects of FAP proliferation and cell-cycle exit that are not fully captured by ligand loss alone. This could include GLI-independent SMO outputs, such as Gi-dependent changes in cAMP signaling or RhoA/ROCK-dependent cytoskeletal regulation (Teperino et al. 2014, Ho Wei et al. 2018, Chechekhin et al. 2022). Consistent with this idea, FAP^no SMO^ muscles show persistent FAP proliferation and elevated FAP numbers at 7 dpi (Fig. 3B). This prolonged FAP expansion could provide a cellular explanation for the increased collagen deposition seen at later stages of repair. In this model, disrupted Hh-dependent FAP cell-cycle exit may lead to FAP accumulation and increased fibrogenic commitment, similar to prior work linking persistent FAP activity to fibrosis (Lemos et al. 2015, Fiore et al. 2016). Future experiments using lineage tracing and SMO mutants that separate canonical GLI activation from non-canonical SMO outputs (Ho Wei et al. 2018, Steiner et al. 2026) will be needed to distinguish these possibilities. For now, the consistent reduction of *Gli1, Timp3*, and *Gdf10* supports impaired canonical Hh signaling as the major driver of the IMAT and regeneration phenotypes, while the fibrosis phenotype may reflect an additional role for SMO in controlling FAP proliferation or fibrogenic remodeling.

### Therapeutic implications of FAP-intrinsic Hh signaling competence for Hh manipulation

The failure of SAG to rescue the FAP^no SMO^ phenotype has important implications for Hh-based therapeutic strategies. It shows that systemic Hh activation only works if FAPs are still able to respond through SMO. When SMO is removed from FAPs, SAG no longer suppresses adipocyte accumulation and no longer improves myofiber regeneration, even though other cell types in the muscle may still be Hh competent. This suggests that FAPs are not just one possible target of Hh signaling, but the key cellular target needed for its regenerative effects. At the same time, the reduced myofiber size seen in SAG-treated control muscles reinforces an important limitation. Hh activation is not always beneficial. As we have shown here and before, its effects depend on timing, dose, and cellular context (Fig. 5) (Kopinke et al. 2017, Norris et al. 2023). Thus, therapeutic strategies should not simply aim to activate Hh signaling broadly throughout the tissue. Instead, they will need to restore or maintain the right level of Hh signaling competence in FAPs. This may be especially relevant in aged and dystrophic muscle, where Hh activity is reduced (Renault et al. 2013, Piccioni et al. 2014, Norris et al. 2023), raising the possibility that inadequate FAP Hh signaling is a shared driver of pathological remodeling across multiple disease settings.

Taken together, this study provides direct genetic evidence that FAPs are the primary cellular mediators of Hh signaling in injured muscle. FAP-intrinsic Hh signaling suppresses IMAT, limits fibrotic remodeling, and maintains the pro-myogenic FAP secretome needed to support muscle stem cell-driven repair. By showing that pharmacological Hh activation fails when FAPs cannot respond, our findings resolve the central question of which cell type functions as the Hh sensor during muscle regeneration. They also point to FAP Hh signaling as a key pathway to preserve or restore in diseases marked by failed regeneration, fatty infiltration, and fibrosis.

## Resource availability

### Lead contact

Requests for further information and resources should be directed to and will be fulfilled by the lead contact, Daniel Kopinke.

### Materials availability

This study did not generate new unique reagents. Mouse lines used are available from The Jackson Laboratory as indicated in the methods.

### Data and code availability

- All data reported in this paper will be shared by the lead contact upon reasonable request.
- No original code was generated in this study. Image segmentation used the published LabelsToROIs pipeline (Waisman et al., 2021; https://github.com/morriscr/LabelsToROI).
- Any additional information required to reanalyze the data reported in this paper is available from the lead contact upon request.

## Acknowledgement

We thank the members of the Kopinke laboratory for critical discussions. This work was supported by the US National Institutes of Health (NIH) grant 1R01AR079449 and 1R01HL171050 to D.K., and the UF Thomas Maren Junior Research Excellence Fund to D.K. The content is solely the responsibility of the authors and does not necessarily represent the official views of the NIH. This work is, in part, the result of NIH funding and is subject to the NIH Public Access Policy. All schematic figure models were created with BioRender.

## Author contributions

Conceptualization, X.L. and D.K.; methodology, X.L. and D.K.; investigation, X.L, V.H., B.D., A.N; writing –original draft, X.L. and D.K.; writing – review & editing, X.L. and D.K.; funding acquisition, D.K.; supervision, D.K. All authors discussed the results and commented on the manuscript.

## Methods

### Mouse lines and husbandry

All animal procedures were approved by the IACUC of the University of Florida and conducted in accordance with the NIH Guide for the Care and Use of Laboratory Animals. All mice were maintained on a 129S/J background under a 12-hour light/dark cycle with ad libitum access to food and water. FAP-specific conditional *Smo* knockout mice (FAP^no SMO^) were generated by crossing *Pdgfrα*^*CreERT*^ mice (Chung et al. 2018) (JAX# 032770) with *Smo*^fl/fl^ mice (Long et al. 2001) (JAX# 004526). For lineage tracing, the Rosa26-EYFP allele (Srinivas et al. 2001) (JAX# 006148) was introduced into the FAP^no SMO^ background. Control mice were littermates lacking the *Pdgfrα*-Cre transgene or carrying wild-type *Smo* alleles. Both male and female mice aged 8–12 weeks were used. Male and female data were pooled for main figure analyses as described below.

### Cardiotoxin-induced muscle injury

Acute skeletal muscle injury was induced by intramuscular injection of cardiotoxin (CTX; Latoxan, *Naja pallida*; Cat# L8102-1MG) as previously described (Kopinke et al. 2017, Norris et al. 2023). CTX was dissolved in sterile saline to 10 µM and 30-50 µL was injected per TA under isoflurane anesthesia. Mice were euthanized by isoflurane inhalation at 3, 7, or 21 dpi as specified.

### Smoothened agonist (SAG) treatment

For SAG 21k treatment (Tocris; Cat# 5282) a stock solution was made in DMSO. This stock was further diluted in sterile PBS and administered at 2.5 mg/kg through IP injection on day 0-, 2- and 4 dpi, as previously described (Norris et al. 2023). Investigators performing quantification were blinded to treatment group.

### Tissue processing and cryosectioning

Tissue processing was performed as previously described (Kopinke et al. 2017, Norris et al. 2023). TAs were fixed in 4% PFA for 2.5 hours at 4°C, washed and cryoprotected in 30% sucrose overnight at 4°C, embedded in OCT-filled cryomolds (Sakura Finetek; Cat# 4566), and snap-frozen in liquid N_2_-cooled isopentane. Cryosections of 12 µm thickness were collected every 250-350 µm using a Leica cryostat.

### Immunofluorescence staining

Immunofluorescence was performed as previously described (Kopinke et al. 2017, Norris et al. 2023). Sections were blocked in blocking solution (5% donkey solution in PBS with 0.3% Triton X-100) at 4°C for 45 minutes at room temperature. Primary antibodies in blocking solution were applied overnight at 4°C: anti-PDGFRα (R&D Systems; Cat# AF1062; 1:250), anti-ARL13B (Proteintech; Cat# 17711-1-AP; 1:1000), anti-SMO (7TM; Cat# 7TM0239A-IC; 1:1000), anti-PERILIPIN (Cell Signaling Technology; Cat# 9349; 1:1000), anti-GFP (Aves Labs; Cat# 1020; 1:500). Species-appropriate Alexa Fluor secondary antibodies (Thermo Fisher Scientific; 1:1000), DAPI (1 µg/mL), and Phalloidin-Alexa Fluor 647 (Thermo Fisher Scientific; Cat# A22287; 1:500) in blocking solution were applied for 1 hour at room temperature. Slides were mounted with Fluoromount-G (SouthernBiotech; Cat# 0100-01). Images were acquired on a Leica DMi8 using a 20x objective. Images of the whole TA were obtained with the navigator function within the Leica LSA software. All images were quantified through ImageJ Software (v1.552p).

### SMO ciliary colocalization quantification

Confocal image stacks (40× oil, NA 1.4) were acquired as previously described (Kopinke et al. 2017). ARL13B^+^/SMO^+^ cilia colocalization was scored per PDGFRα^+^ FAP using a fixed fluorescence threshold applied uniformly across images. A minimum of 5 fields per TA were averaged for per-mouse statistical analysis.

### Sirius red staining and fibrosis quantification

Sirius red staining was performed as previously described (Norris et al. 2023). Fibrotic area was quantified in FIJI/ImageJ using a fixed HSB hue threshold and expressed as a percentage of total injured TA area (Norris et al. 2023).

### Myofiber cross-sectional area quantification

Myofiber CSA was quantified using the Cellpose segmentation algorithm and LabelsToROIs FIJI/ImageJ plugin as previously described and validated (Waisman et al. 2021, Norris et al. 2023). Whole TA tissue section were segmented, eroded 1 pixels to correct for PFA fixation halo, and individual fiber ROI measurements exported as CSV. Myofibers with CSA < 50 µm^2^ were excluded. Distributions were binned in 600 µm^2^ increments. Myofibers were pseudo-color coded by CSA using the ROI Color Coder macro.

### Intramuscular adipocyte quantification

PERILIPIN^+^ adipocytes were manually counted from tile-scan images of complete TA cross sections as previously described (Waisman et al. 2021, Johnson et al. 2022, Norris et al. 2023), and normalized to injured area (mm^2^). For lineage tracing, PERILIPIN^+^/GFP^+^ cells were expressed as a fraction of total PERILIPIN^+^ adipocytes. All counting was performed blinded to genotype and treatment group.

### RNA extraction and RT-qPCR

RNA extraction and RT-qPCR were performed as previously described (Kopinke et al. 2017, Norris et al. 2023). Total RNA was extracted using TRIzol reagent (Thermo Fisher Scientific; Cat# 15596026). cDNA was synthesized from 500 ng RNA using the qScript Reverse Transcriptase kit (Quanta Bio; Cat# 84003). RT-qPCR was performed on a QuantStudio 6 Flex Real-Time 384-well PCR System (Applied Biosystems; Cat# 4485694) using PowerUp SYBR Green Master Mix (ThermoFisher Scientific; Cat# A25742). Expression was normalized to the geometric mean of *Hprt, Sra1*, and *Rpl32* using the 2^−ΔΔCt^ method. Primer sequences are listed in the Supplement.

### Quantification and statistical analysis

TAs with injury less than 50% of total area were excluded from this study. The experimenter was blinded until data was collected. All quantitative data are presented as mean ± SEM and were graphed using GraphPad Prism (version 10). For comparing two samples with one variable, an unpaired two-tailed Student’s t-test was used. For more than two samples with one variable, a one-way ANOVA followed by a Dunnett’s multiple comparison test was used. For two variables, a two-way ANOVA followed by Tukey’s multiple comparison was carried out. Myofiber size frequency distributions were compared by two-way ANOVA with repeated measures across bins; interaction P-values are reported on distribution plots. Sexes were pooled for main analyses with sex indicated by distinct symbols. A p value less than 0.05 was considered statistically significant and denoted as follows: *p < 0.05; **p < 0.01; ***p < 0.001; ****p < 0.0001. The total number of animals used per quantification is listed in the corresponding figure legends.

## Supplementary information

**Supplementary Figure 1.**
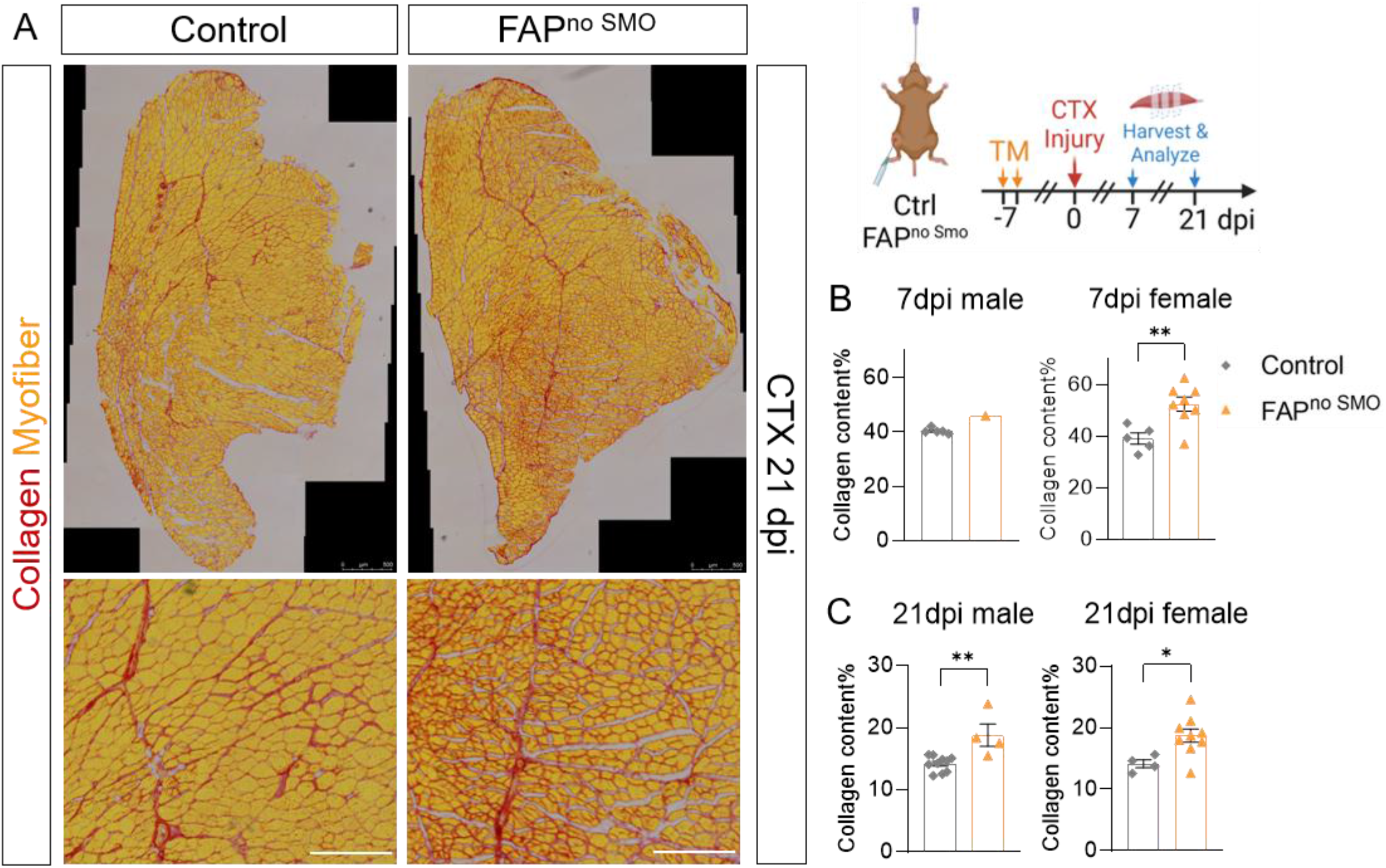
Increased muscle fibrosis in FAP^no SMO^ mice is consistent across sexes and time. **(A)** Genetic scheme and experimental outline, and representative whole-section (scale bar: 500 µm) and zoom-in (scale bar: 300 µm) Sirius red images of TA cross sections from control and FAP^no SMO^ mice at 21 dpi CTX. Collagen, red; myofibers, yellow. **(B)** Quantification of fibrotic area (percentage of injured area) at 7 dpi CTX in (left) male (n = 4–5 mice per group) and (right) female (n = 5–7 mice per group) animals. **(C)** Quantification of fibrotic area at 21 dpi CTX in (left) male (n = 4–9 mice per group) and (right) female (n = 5 mice per group) animals. Data are presented as mean ± SEM. **p < 0.01; *p < 0.05. Statistical comparisons were made by unpaired two-tailed Student’s t-test.

**Supplementary Figure 2.**
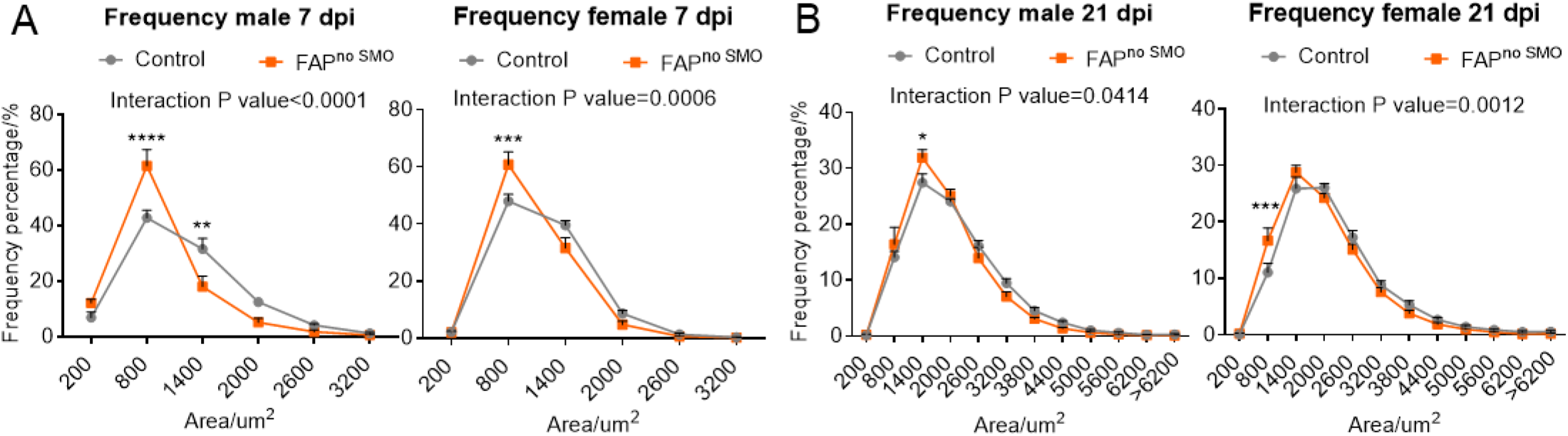
FAPs lacking SMO result in smaller regenerated myofibers *in vivo*. Myofiber size frequency distributions at **(A)** 7 dpi (n = 5–8 mice per sex per group) and **(B)** 21 dpi (n = 5–8 mice per sex per group), shown separately for male and female mice; interaction P values from two-way ANOVA are indicated. Data are presented as mean ± SEM. ****p < 0.0001; ***p < 0.001; **p < 0.01; *p < 0.05.

**Supplementary Table 1.**
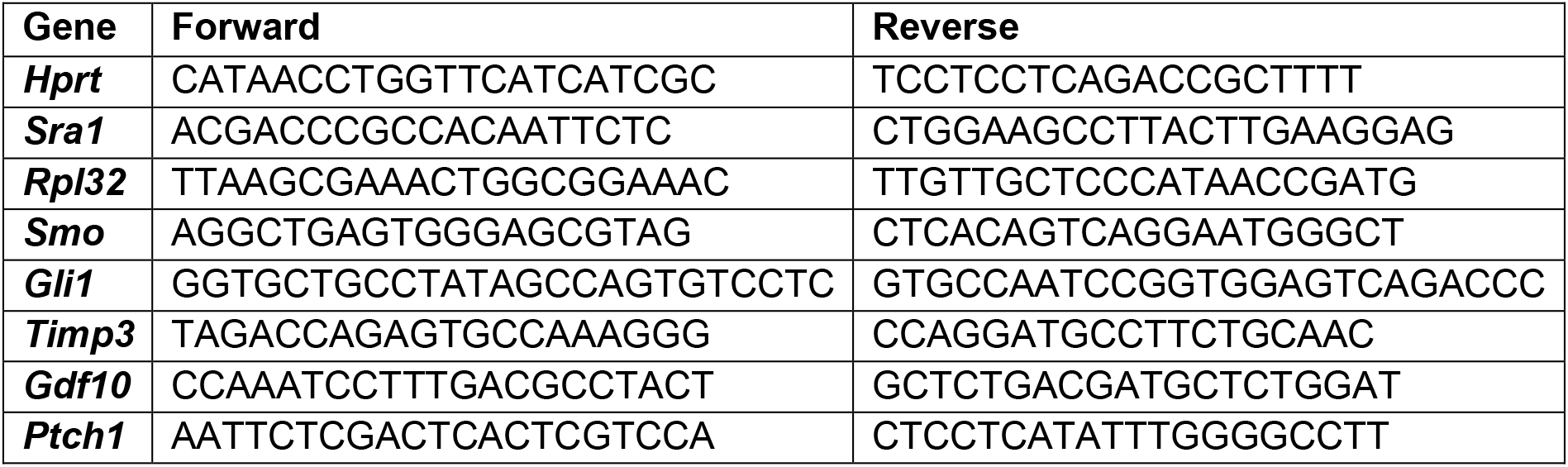
The qPCR primers used in the experiments.

